# From cyst to tapeworm: Transcriptome of *Taenia solium in vitro* activation and growth

**DOI:** 10.1101/2025.02.04.636528

**Authors:** David Castaneda-Carpio, Renzo Gutierrez-Loli, Jose Maravi-Jaime, Segundo W Del Aguila, Valeria Villar-Davila, Luz M Moyano, Rafael Tapia-Limonchi, Stella M Chenet, Cristina Guerra-Giraldez

**Affiliations:** Laboratorio de Proliferación Celular y Regeneración, Facultad de Ciencias e Ingeniería, Universidad Peruana Cayetano Heredia. Lima, Peru; Centro de Salud Global, Tumbes, Universidad Peruana Cayetano Heredia, Peru; Escuela Profesional de Medicina Humana, Universidad Nacional de Tumbes. Tumbes, Peru; Instituto de Investigación de Enfermedades Tropicales, Universidad Nacional Toribio Rodríguez de Mendoza de Amazonas. Chachapoyas, Peru; Facultad de Medicina, Universidad Nacional Toribio Rodríguez de Mendoza de Amazonas. Chachapoyas, Peru

## Abstract

The cestode *Taenia solium* develops as a tapeworm solely in the human intestine, starting from a larva (cyst). Upon maturing, it produces hundreds of thousands of infectious eggs. When ingested by pigs or humans, the eggs develop as cysts that lodge in various tissues, including the brain, leading to neurocysticercosis. Despite advances in understanding cestode biology through genomic and transcriptomic studies, particularly in model organisms, much remains unknown about the activation of *T. solium* cysts in the human digestive tract and the events that drive the development into adult worms—the stage responsible for dispersing the parasite. We present a transcriptome generated by Next Generation Sequencing from *T. solium* cysts activated in culture and collected at three different in vitro growth phases, as defined by their morphology. We identified differentially expressed genes and biological processes relevant to growth phases, which can be further explored with the dataset. The information is valuable for identifying genes that regulate the molecular, metabolic, and cellular events leading to parasite maturation or elements driving its transmission.

## Background and Summary

*Taenia solium* is a parasitic flatworm causing neurocysticercosis and taeniasis. Neurocysticercosis, from ingesting *T. solium* eggs, leads to neurological symptoms and can be fatal. This neglected tropical disease is common in areas with poor sanitation and free-roaming pigs, causing up to 70% of epilepsy cases where it is endemic (1). The World Health Organization (WHO) identifies *T. solium* as a major cause of death from food-borne diseases, with a 2.8 million disability-adjusted life-years (DALYs) globally (2). Infected animals also cause economic losses for farmers (3).

*T. solium*’s life cycle involves pigs as intermediate hosts and humans as the only definitive hosts. The adult tapeworm sheds embryonated eggs in human feces. When ingested by pigs or humans, embryos travel mainly to muscles, the heart, and the brain while developing into vesicular cysts (4). These cysts, if consumed in undercooked pork, are activated by digestive enzymes and bile salts. The activation triggers the evagination of the parasite’s scolex, a process where the scolex, containing hooks and suckers, emerges from the cyst (5). The evagination is driven by cell proliferation at the neck, where totipotent cells play a crucial role. These cells facilitate the eversion of the scolex, which can then attach to the human intestine. (6). The parasite grows continuously by generating segments (proglottids) that produce eggs, perpetuating the cycle (4,7,8).

Humans thus carry the parasite’s dispersal form, the tapeworm with mature proglottids. A few carriers can infect many pigs in the rural context of a developing country (9,10). One Health strategies aim to control the disease by treating pigs and humans (11). Mathematical models like CystiAgent highlight the importance of reducing the tapeworm’s infectious duration to curb human-to-porcine transmission (9,12). However, the biology of this stage is the least known, and studying it is challenging, due to ethical and practical issues associated with humans as hosts.

The parasite’s genome (13) and partial transcriptomes have provided insights into their development (14,15). Comparative studies, including whole-transcriptome analyses with Next Generation Sequencing of cestodes *Echinococcus* (16), *Hymenolepis* (17), and other *Taenia* (18,19), have identified common genes and pathways. Still, the life cycles of these species differ significantly from *T. solium*, leaving a critical gap in understanding *T. solium*’s specific molecular and cellular mechanisms during cyst activation and subsequent development into the adult worm. Single-gene approaches have focused on the early development of bile-activated cysts (20). Bile salts have long been used to catalyze the *in vitro* transition to the adult stage (21,22), but the transcriptomic program triggered by this treatment—mimicking cyst activation in the human intestine—has not been characterized.

Our dataset provides information about *T. solium*’s cyst activation and growth into the adult intestinal form. We incubated fresh cysts extracted from naturally infected pig muscle, adding taurocholic acid (TA) for cyst activation. We determined three morphological phases during scolex evagination, collected samples at each phase, and performed RNA-Seq to compare their transcriptomic profiles. We assessed the sequencing reads quality, mapped them to the *T. solium* reference genome, and performed differential gene expression analysis to compare their transcriptomic profiles and identify differentially expressed genes. (Fig. 1a)

**Figure 1.**
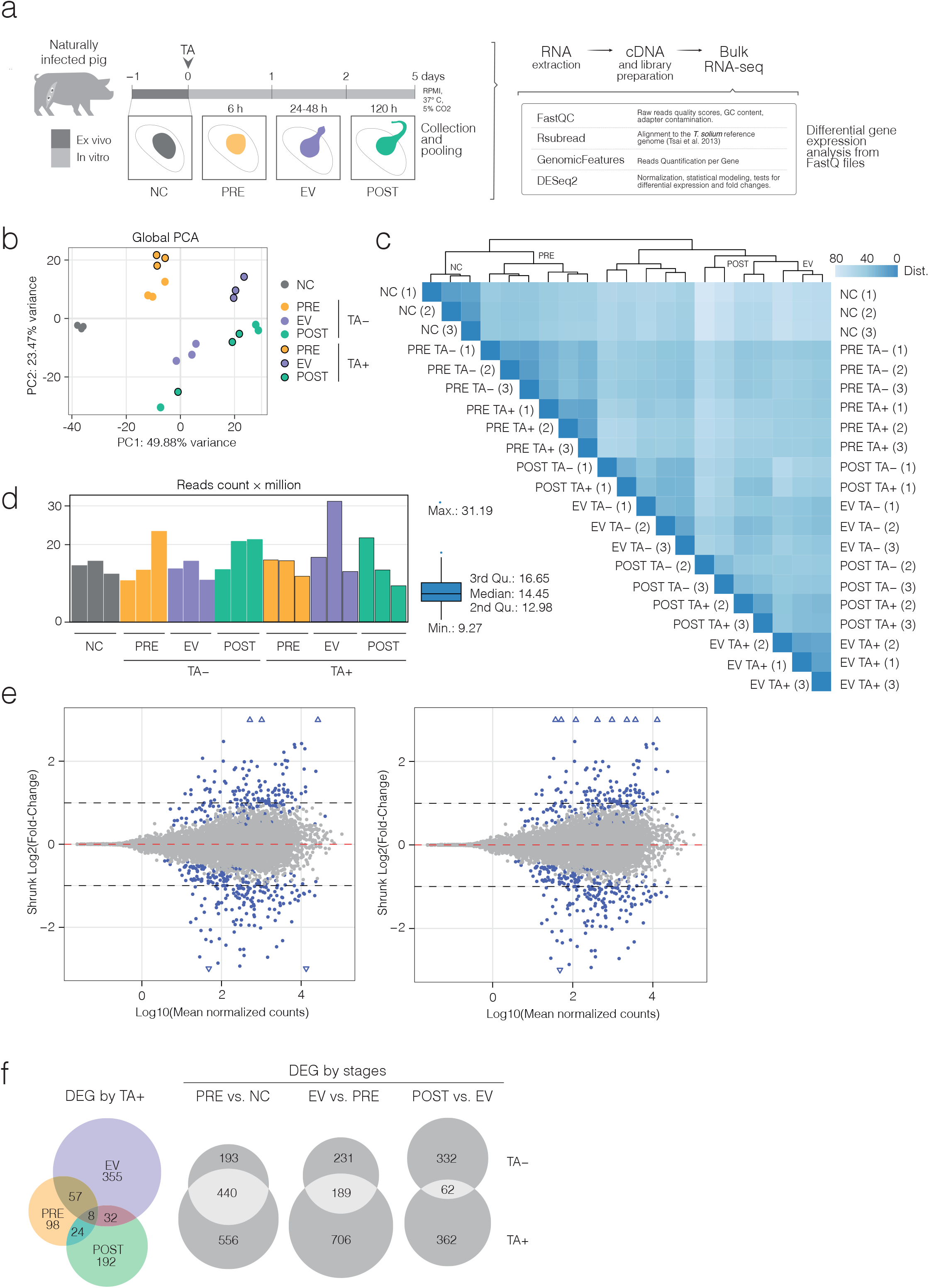
Study design for in vitro evagination and Technical Validation of the RNA-seq analysis by parasite phase and TA treatment. (a) *T. solium* cysts were cultured ex vivo for up to 5 days and stimulated with TA. Phases were determined based on macroscopic morphological differences. Whole RNA was extracted from pooled parasites at each phase (NC, PRE, EV, POST), followed by cDNA library preparation and sequencing using Illumina’s NextSeq 500 platform. (b) Principal Component Analysis (PCA) of the 500 most variable genes, determined by the coefficient of variation (CoV), reveals clustering by parasite group and phase (represented by colors) and TA treatment (denoted by outlined circles). (c) Hierarchical clustering of samples, based on Euclidean distance using the CoV of all the geneset, underscores phase-specific and treatment-related groupings. (d) Bar plot showing the number of mapped reads per replicate across NC, PRE, EV, and POST cysts, with or without TA treatment. The accompanying boxplot represents the minimum and maximum values, median, and interquartile range (2nd and 3rd quartiles). (e) MA plots displaying the shrunk log_2_ fold-change relative to the log_10_ of mean normalized counts for the comparison between EV vs. PRE, in the absence (left) or presence (right) of TA. (f) Euler plots illustrate differentially expressed genes (DEGs), induced or repressed by TA across PRE, EV, and POST parasites. Additionally, the plots compare DEGs between phases (PRE vs. NC, EV vs. PRE, and POST vs. EV) in the presence or absence of TA.

The dataset can serve as a starting point for exploring condition-specific gene expression patterns. The groups selected for the transcriptome allow to discriminate between events happening on parasites that had recently evaginated the scolex and were induced by TA (EV TA+), and those that had not (PRE TA+). On these groups, TA induction affected 895 transcripts; 403 were upregulated, while 492 were downregulated (Fig. 2a). Specifically, 309 transcripts were upregulated and 398 downregulated after TA-induced evagination of the scolex.

**Figure 2.**
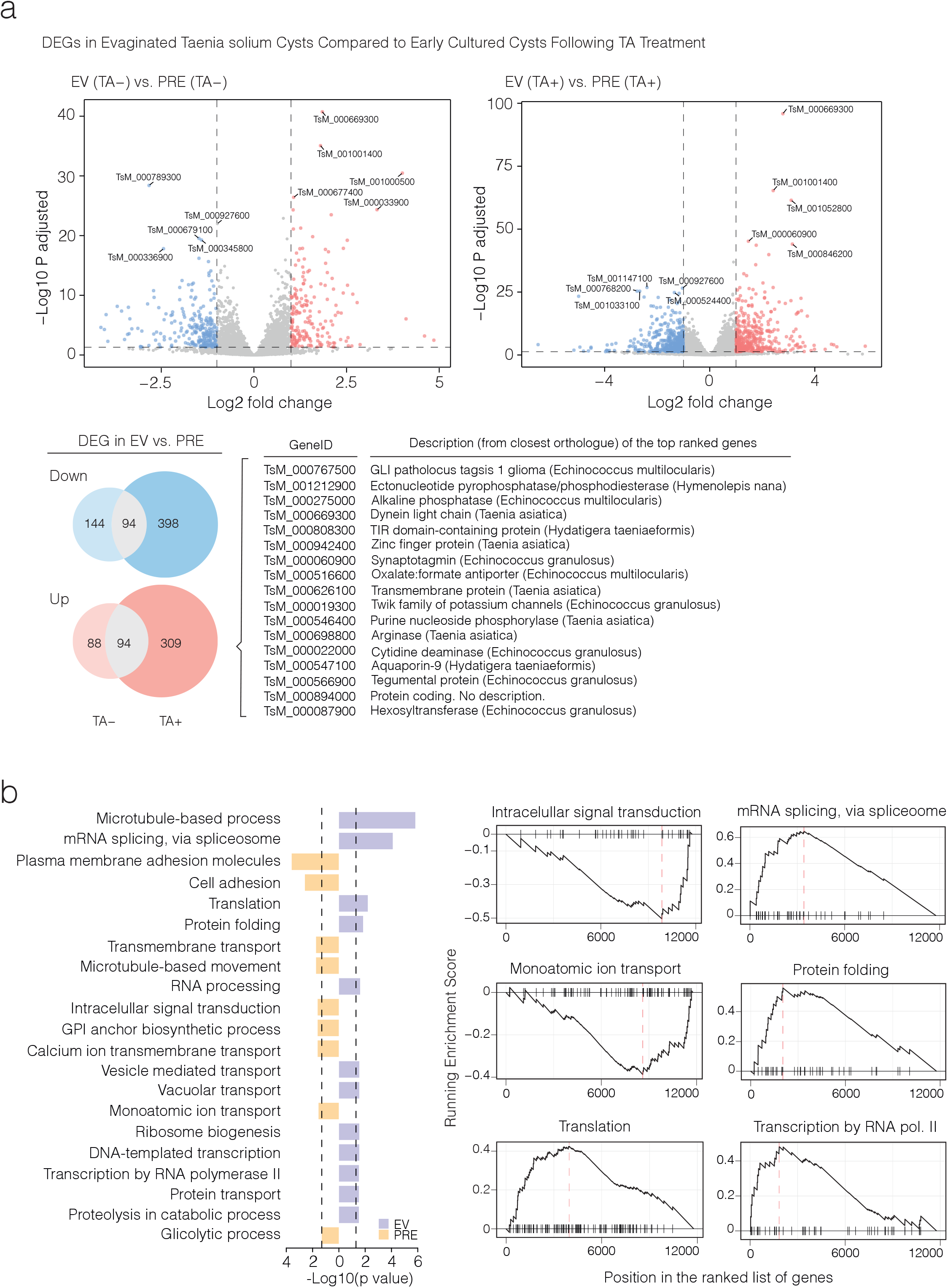
Visualization and exploration of phase- and TA-specific RNA-seq data of *T. solium* cultured cysts: Informative comparisons. (a) Differentially expressed genes (DEGs) in *Taenia solium* EV cysts compared to PRE cysts following TA addition. The upper panel shows volcano plots of DEGs comparing EV and PRE cysts according to TA’s absence (left) or presence (right). Each dot represents a gene, with red indicating significantly upregulated genes and blue, significantly downregulated genes (adjusted p < 0.05). Labeled genes correspond to top-ranked DEGs based on significance. The lower panel presents Euler diagrams illustrating the overlap of DEGs (upregulated and downregulated) between EV and PRE cysts under TA-free (TA−) and TA-treated (TA+) conditions, along with a table listing the top-ranked DEGs, their gene IDs, and descriptions from the closest orthologous genes. (b) Gene ontology enrichment and gene set enrichment analysis (GSEA) of differentially expressed genes in evaginated *Taenia solium* cysts. The left panel presents a bar plot of significantly enriched Gene Ontology (GO) biological processes for differentially expressed genes (DEGs). The x-axis represents the −log_10_(*p*-value), with higher values indicating greater statistical significance. Bars are color-coded to distinguish between EV-related (yellow) and PRE-related (purple) processes. The right panel displays gene set enrichment analysis (GSEA) plots for selected biological processes. The running enrichment score is shown for each gene set, with the red vertical dashed line indicating the position of the leading-edge subset, which contributes most to the enrichment signal.

Few *Taenia solium* genomes and transcriptomes are available; therefore, reliable functional annotations for each gene are lacking. In the WormBase ParaSite database only 6,168 (49.5%) have at least one associated GO term, with a mean of 2.499 GO terms per gene. To provide more elements to analyze and compare, we enhanced the data available in public repositories by functional annotation with blast2GO (23). This added 3,317 newly annotated genes (26.6% of the total 12,467 *T. solium* coding sequences (13)) and 9,821 extra Gene Ontology (GO) terms. Our blast2GO annotation assigned at least one GO term to 9,485 (76.1%) genes, with a mean of 15.426 GO terms per gene, leaving only 2,982 (23.9%) genes without annotation. Additionally, we generated a custom annotation package with AnnotationForge (24) to facilitate over-representation analysis using clusterProfiler (25), enabling a more detailed exploration of the biological processes active in our dataset.

Further exploration by GSEA of EV and PRE TA+ shows that transcripts of the functional categories “Monatomic ion transport” and “Intracellular signal transduction” were more active before evagination of the scolex (PRE TA+), and “Transcription by RNA polymerase II”, “mRNA splicing via spliceosome”, “Translation” and “Protein folding” were enriched after evagination of the scolex (EV TA+) (Fig. 2b).

This dataset offers the first transcriptome focused on the *T. solium* cyst activation and development into a tapeworm, the stage naturally occurring inside the human host. The data also includes new annotations for 76.1% of the coding genes present in WormBase ParaSite. By associating transcripts to phases surrounding cyst activation, the data provides a foundation for future research on gene function in *T. solium* early-adult development and a model to examine scolex evagination and strobilation at multiple molecular, cellular, histological, and morphological levels. Further investigation could uncover regulatory elements and pathways triggering and modulating the maturation of *T. solium* in the human intestine.

## Methods

### Ethics statement

The study protocol was revised and approved by the Animal Care and Use Committee (CIEA, in Spanish) at Universidad Peruana Cayetano Heredia (Certificates CIEA 040-09-22, R 040-09-23 and R 040-09-24, valid until August 2025). The CIEA states details for the proper housing, feeding, and handling of animals used in research, including euthanasia. This Committee is registered with the Office of Laboratory Animal Welfare of the Department of Health and Human Services of the National Institutes of Health (NIH – USA) with Assurance Number F16-00076 (A5146-01), valid until August 31, 2026.

### Collection of *T. solium* cysts from pig muscle

*Taenia solium* cysts (n=500) were collected from the muscle tissue of a naturally infected pig at Universidad Peruana Cayetano Heredia’s Global Health Center in Tumbes, Peru. Upon collection, cysts were washed three times with transport solution (1X PBS, pH 7.2, with Gibco™ Antibiotic-Antimycotic to the following final concentrations: 100 units/mL penicillin, 100 μg/mL streptomycin and 250 ng/mL amphotericin B). Cysts were placed in 50-mL sterile Falcon tubes (up to 100 cysts per tube) with fresh transport solution and packed to be delivered by air freight to the university laboratories in Lima.

### *In vitro* scolex evagination and morphology of activated parasites

Cysts arrived at the laboratory approximately 16 hours after collection and were washed three times with transport solution. After shipment, most cysts maintained their characteristics: intact, white, or translucent oblong-shaped vesicles without atypical coloration, filled with a clear fluid. Cysts were then distributed for different purposes; 84 were used for the present study.

Twelve cysts were placed in 1 mL of RNAlater® solution (R-0901, Sigma Aldrich) and stored at - 20°C until RNA extraction and gene transcription analysis. These *ex vivo* cysts were defined as non-cultured (NC).

Each one of 72 cysts was incubated in 1.5 ml filtered sterilized RPMI 1640 medium (pH 7.4), supplemented with 2 g/mL NaHCO_3_ (Merck®, Burlington, MA), 1.6 µM β-mercaptoethanol, and Gibco™ Antibiotic-Antimycotic in the concentrations described above. One cyst per well was placed in three Costar® 24-well clear-tissue Culture-treated surface plates (Corning®, Corning, NY, USA) at 37°C and 5% CO_2_ in a sterile incubator (VWR Symphony Incubator) for up to five days. The medium of 36 of the 72 cysts included 0.1% taurocholic acid (TA) (Sigma, St. Louis, MO, USA). Therefore, half of the samples were TA+ and half TA-. The medium was replaced every 24 hours. Cysts were collected in three distinct moments: cysts with non evaginated scolex, after 6 hours *in vitro*, TA+ and TA-(defined as “pre-evagination”: PRE TA+ and PRE TA-); cysts with recently exposed scolex, approximately after 24-48 hours *in vitro* (defined as “recent evagination”: EV TA+ and EV TA-); and cysts having fully evaginated, already showing some proglottids, after 48-120 hours *in vitro* (defined as “post-evagination”: POST TA+ and POST TA-). After culture, all parasites were washed with 1X PBS to remove residual media, and preserved at -20°C in 1 mL RNAlater® solution until RNA extraction and gene transcription analysis. Evagination curves were made along the 240 hours *in vitro*.

### RNA isolation

RNA was extracted from 12 samples (cysts) of each of the seven groups (NC, PRE TA+, PRE TA-, EV TA+, EV TA-, POST TA+, POST TA-; n=84 cysts) using the Quick-RNA Miniprep Plus Kit (Zymo Research, R1058). Briefly, samples were independently embedded in 600 µL of DNA/RNA Shield (Zymo Research, R1058) and admixed with a tissue homogenizer tip (Omni International, Kennesaw, GA) on ice, followed immediately by a 30-min treatment with 60 mg of Proteinase K (Zymo Research, D3001-2-60). RNA was isolated following the manufacturer’s protocol, with an extra washing step with 400 µL of RNA Wash Buffer (Zymo Research, R1058) before final RNA isolation. The protocol included DNAse I treatment (Zymo Research, E1009-A (250 U)). As recommended by the manufacturer, a mix of 5 µL of DNase I (1 U/µL) and 75 µL of DNA Digestion Buffer (Zymo Research, R1058) was added directly to the silica column matrix (Zymo-Spin™ IIICG Columns) containing processed samples, for a 15-min incubation at room temperature (20-30°C). RNA yield was measured by spectrophotometry with NanoDrop® (Thermo Fisher Scientific, Waltham, MA, USA), and approximately 300 ng of RNA were loaded into a 1.5% agarose/1% bleach gel to test its integrity by electrophoresis (35 minutes at 100V) (26)

### RNA-Seq library construction and sequencing

We constructed 21 libraries, three for each of the seven groups (NC, PRE TA+, PRE TA-, EV TA+, EV TA-, POST TA+, POST TA-). Each library was made from four cysts, pooling 125 ng of independently extracted RNA for a final 500 ng/library. Following the manufacturer’s instructions, the Illumina® Stranded mRNA Prep Ligation Kit (Illumina, 20040532) was used to construct libraries indexed with the IDT for Illumina RNA Unique dual (UD) Indexes. Briefly, Oligo(dT) RNA purification magnetic beads (RPBX, Illumina, 20040532) allowed for polyA-RNA selection and excluded ribosomal RNA. Fragmentation of purified mRNA, cDNA synthesis of the first and second strands, and ligation of the Illumina RNA UD Indexes Set A followed. The libraries were amplified by 13 PCR cycles and quantified using the Qubit^TM^ dsDNA High Sensitivity (HS) Assay Kit (Thermo Fisher Scientific, Q32851). The library fragment sizes and their integrities were verified on 2% agarose gel electrophoresis.

Each library was diluted to a starting concentration of 1nM, and the 21 libraries were combined using 10 μL of each diluted library. The combined libraries were denatured and diluted to a final concentration of 1.4 pM before sequencing. Libraries were sequenced for 75 cycles in paired-end mode with a high output kit on the Illumina NextSeq 500 platform (Illumina, San Diego, CA, USA) according to the Illumina protocol. FASTQ files were generated using the bcl2fastq software (Bcl2Fastq; Illumina). FASTQ data was deposited in GEO (GSE288552).

### RNA-seq data analysis

Quality control of the reads was assessed with FastQC (version 0.12.0) (27). Removal of Illumina adapter sequences and trimming of the reads with a phred+33 quality score below 20 was done using BBDuk (BBMap version 38.90) (28). Reads were mapped to the *T. solium* genome (assembly GCA_001870725.1) (13) using the default settings of RSubread (version 2.17.4) (29). Gene abundance was calculated with GenomicFeatures (v. 1.52.1) (30) using predicted gene annotations available in the WormBase ParaSite database (version WBPS18) (13,31).

Differential expression analysis was performed using a generalized linear model that assumes a negative binomial distribution of count data, implemented by DESeq2 (v.1.40.2) (32). Differentially Expressed Genes (DEGs) were defined as such when their mean expression differed between conditions by at least 2-fold and the calculated False Discovery Rate (FDR) adjusted p-values by the Benjamini-Hochberg procedure were equal to or less than 0.05.

DESeq2 variance-stabilized counts were used for a principal component analysis (PCA) using the 500 genes with the highest variance. Additionally, hierarchical clustering was performed across all sample counts by calculating an Euclidean distance-based matrix.

### Functional annotation of the *T. solium* genome

We retrieved full-length *T. solium* transcript sequences from the WormBase Parasite database (PRJNA170813) (13,31). After importing the sequences into blast2GO (OmicsBox suite) (23), we replaced duplicate IDs and added descriptions. A DIAMOND blastx search was conducted against the NR database with an E-value of 1×10^-5^ and a taxonomy filter for Platyhelminthes (6157), Trematoda (6178), Cestoda (6199), and Nematoda (6231).

We used InterProScan to identify conserved domains, motifs, and signal peptides, applying various databases (e.g., Pfam, Panther, SMART, SignalP). High-confidence functional annotations were assigned using an E-value threshold of 1×10^-6^, an annotation cutoff of 55, and a GO weight of 5. Additionally, we integrated GO annotations from eggNOG 5 (33) to enhance annotation quality with orthologous group information using a seed ortholog E-value filter of 1×10^−5^ and a seed ortholog bit-score filter of 60.0.

Finally, we exported the annotated data into R, creating a custom annotation package using the makeOrgPackage() function from AnnotationForge (24) for downstream analyses. This process provides a solid framework for exploring the functional roles of *T. solium* genes across different conditions.

### Over-representation Analysis (ORA) of functional categories

Gene Set Enrichment Analysis (GSEA) was performed using the gseGO() function (eps = 1×10^-300^, ont = “BP”) from the clusterProfiler package (25). We ranked genes in decreasing order based on the Wald statistic obtained from differential gene expression analysis with DESeq2. The 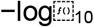 of the corrected p-values were plotted for all enriched GO terms with a threshold of 0.05. GSEA plots for each functional category were generated using the gseaplot() command from clusterProfiler.

### Data Records

This dataset consists of bulk RNA-seq data from 21 samples, representing seven experimental groups, each with three biological replicates. Each sample comprises four pooled *Taenia solium* cysts. The groups include non-cultured parasites (NC); non-evaginated parasites cultured for 6 h without TA (PRE TA-) or with TA (PRE TA+); early-evaginated parasites cultured between 24 and 48 h without TA (EV TA-) or with TA (EV TA+); and fully evaginated parasites cultured for 120 h without TA (POST TA-) or with TA (POST TA+). The raw sequencing data and gene counts matrix are available in the Gene Expression Omnibus (GEO) under accession number GSE288552, while the *T. solium* genome can be accessed via WormBase ParaSite (13,31).

Our GitHub repository (https://github.com/dcastanedac/ts-seq) contains scripts for validating and analyzing the transcriptome of *T. solium* activated cysts, which will allow the users interested in the data set to explore and mine data to generate hypotheses. The pipeline includes quality control, trimming, read mapping, and count matrix generation for validating read quality and mapping to the *T. solium* reference genome. It also features clustering analysis for experimental group validation, filtering and visualization of differentially expressed genes (DEGs), and functional annotation using over-representation analysis (ORA). Data analysis scripts cover tasks such as PCA, hierarchical clustering, DEG comparison, GSEA, and functional annotation, with corresponding GO annotations and files for generating a custom annotation package. In addition, our repository contains raw count matrices (counts.RData), the DESeq2 object (dds.RData) and GO annotations (Tsolium_annotation_full-export.xlsx).

### Technical Validation

All cultivated cysts (elongated vesicles, approximately 1 cm long) looked healthy, with a smooth surface and clear content. When measured by NanoDrop, the A260/A280 ratios for RNA were between 1.8-2.0. Additionally, the 18S and 28S ribosomal RNA bands were clearly visible in total RNA run on denaturing 1.5% agarose gel. All 21 libraries run on 2% agarose gel showed a minimum size of ∼ 250 bp in length. We estimated that our 21 libraries could be sequenced with one cartridge using the NextSeq 550 system with the High Output kit v2.5 from Illumina (Illumina, U.S.A.). Based on the human genome (3.1 billion base pairs), the commercial recommendation is 50 million reads for a transcriptome (16 transcriptomes with one cartridge). Considering that the *T. solium* genome (around 122 million base pairs) is 25 times smaller than the human genome, we estimate that 30 million reads are enough to cover each of the transcriptomes we wish to compare. Therefore, 400 million reads allowed sequencing of the seven groups’ biological triplicates (i.e., 19 million reads per transcriptome). These triplicates are also based on pools of individuals, giving more power to the output.

After quality control with FastQC, only 8.60±5.61% of the reads had to be trimmed due to quality adapter sequence contamination and low read quality (phred+33 < 20). An average of 15.89 million mapped reads were counted for each replicate. An average of 15.89 million mapped reads were counted for each replicate. In addition to that, almost all the detected reads (98.47±0.19%) were uniquely mapped to the reported *T. solium* genome (Fig. 1d).

Data replicability was assessed by performing a principal component analysis (PCA) on the counts per sample for the 500 most variable genes (Fig. 1b). The first two principal components (PCs) accounted for 73.35% of the total variance. PCA revealed clustering of biological replicates according to their experimental condition, except for POST parasites in both TA- and TA+, likely reflecting higher variability due to prolonged *in vitro* conditions. Notably, EV parasites, whether stimulated or not with TA, formed two distinct clusters. PRE parasites clustered together and apart from NC, EV, and POST parasites.

Hierarchical clustering of samples (Fig. 1c) further highlighted a single cluster encompassing NC and PRE parasites. PRE parasites formed two subclusters based on treatment, while NC and PRE parasites were clearly distinct in gene expression from EV and POST parasites. Additionally, the impact of TA on gene expression in EV cysts was evident, as they formed two well-differentiated subclusters based on treatment. Within the EV-POST cluster, two replicates of POST TA- and two replicates of POST TA+ grouped with the EV TA+ parasites, while one POST TA- and one POST TA+ replicate clustered with the EV TA- group, pointing at transcriptional heterogeneity among the POST parasites.

MA plots of the shrunk fold changes between non-evaginated and early-evaginated parasites (±TA) show that most of the detected DEGs do not have low count levels (mean normalized counts < 10) which would allow bias in quantification due to high levels of variance (Fig. 1e).

When analyzing differentially expressed genes (DEGs) between developmental phases within each treatment condition (Fig. 1f), we identified 98 DEGs exclusive to PRE, 355 exclusive to EV, and 192 exclusive to POST in the presence of TA. Eight DEGs were shared across all phases, while 57 DEGs were common between EV and PRE, 32 between EV and POST, and 24 between POST and PRE.

Comparing DEGs between treatments within each phase (Fig. 1F), we found that PRE parasites exhibited 996 DEGs in the presence of TA and 633 in its absence when compared to NC parasites. Of these, 556 were exclusive to TA+ and 193 to TA-, with 440 shared DEGs. EV parasites, compared to PRE parasites, displayed 895 DEGs in the presence of TA and 420 in its absence. Among these, 706 were unique to TA+, 231 were unique to TA-, and 189 were shared between treatments. Lastly, POST parasites, compared to EV parasites, showed 424 DEGs in the presence of TA and 394 in its absence, with 362 exclusive to TA+, 332 exclusive to TA-, and only 62 shared DEGs between treatments.

## Code Availability

Scripts and files used for processing, analysis, and visualization of this transcriptome have been deposited in a GitHub repository (https://github.com/dcastanedac/ts-seq).

## Acknowledgments, Author Contributions & Competing Interests

We thank Jhon Zumaeta and Marianella Villegas from the Instituto de Investigación de Enfermedades Tropicales, Universidad Nacional Toribio Rodríguez de Mendoza de Amazonas, as well as Guillermo Trujillo from the NGS Division at GenLab Peru, for their valuable assistance and guidance during library preparation. We also thank Dr. Pablo Tsukayama from the Laboratorio de Genómica Microbiana, Facultad de Ciencias y Filosofía, Universidad Peruana Cayetano Heredia, Lima, Peru, for providing access to the NGS platform required for the processing of the sequencing cartridge used in this study, and the veterinary team at the Centro de Salud Global Tumbes, UPCH, for their expert support in parasite collection. This work was funded by the Peruvian Programa Nacional de Investigación Científica y Estudios Avanzados, Prociencia (PE501079376-2022).

DCC, JMJ, RGL, and CGG conceptualized and designed this work. LMM supervised the acquisition of parasites. DCC, JMJ, and VVD worked with parasite culture and nucleic acids to prepare the libraries and run the NGS protocols. SDA performed the initial quality control assessment and preprocessing of the RNA-seq data. DCC designed the ad hoc analytical framework for differential comparisons and Gene Ontology (GO) term identification. DCC and RGL conducted data exploration, processing, and mining for primary analyses and dataset validation, and generated the figures. CGG drafted the manuscript and wrote it with input from all co-authors (DCC, RGL, JMJ, SDA, and VVD: transcriptome and analyses; LMM: *T. solium* impact on public health; RTL: cellular processes; SMC: libraries and NGS).

DCC, JMJ, and VVD were supported by a grant from the Peruvian Programa Nacional de Investigación Científica y Estudios Avanzados, Prociencia (PE501079376-2022), which also funded this work.

The authors declare no competing interests.

## Notes

### Competing Interest Statement

The authors have declared no competing interest.

